# Evidence for a force favouring GC over AT at short intronic sites in *Drosophila simulans* and *D. melanogaster*

**DOI:** 10.1101/2021.02.16.431542

**Authors:** Ben Jackson, Brian Charlesworth

## Abstract

Population genetics studies often make use of a class of nucleotide site free from selective pressures in order to make inferences about population size changes or natural selection at other sites. If such neutral sites can be identified, they offer the opportunity to avoid any confounding effects of selection. Here we investigate evolution at putatively neutrally evolving short intronic sites in natural populations of *Drosophila melanogaster* and *D. simulans*, in order to understand the properties of spontaneous mutations and the extent of GC-biased gene conversion in these species. Use of data on the genetics of natural populations is advantageous because it integrates information from large numbers of individuals over long timescales. In agreement with direct evidence from observations of spontaneous mutations in *Drosophila*, we find a bias in the spectrum of mutations towards AT basepairs. In addition, we find that this bias is stronger in the *D. melanogaster* lineage than the *D. simulans* lineage. The evidence for GC-biased gene conversion in *Drosophila* has been equivocal. Here we provide evidence for a weak force favouring GC in both species, which is stronger in *D. simulans.* Some homologous short intronic sites have diverged in GC content between the two species, which may have been caused by lineage-specific changes in the extent to which different regions of the genome are subject to a GC (or AT)-favouring force.

## Introduction

Population genetics studies often make use of a class of nucleotide site that is considered to be free from selective pressures, for the purpose of making inferences about the demographic history of a population (e.g., Fagundes et al. 2007; Obbard et al. 2012; Garud et al. 2015). Such sites are also used as a neutral comparator for the purpose of estimating the parameters of natural selection acting on other types of sites, while controlling for other processes that affect the allelic composition of populations, such as population size changes, genetic drift and mutation. For example, the McDonald-Kreitman test and its extensions (McDonald and Kreitman 1991; Fay et al. 2001; Smith and Eyre-Walker 2002; Welch 2006; Messer and Petrov 2013) rely on a class of site that are assumed to evolve neutrally, whose relative levels of between-species divergence and within-species variability are contrasted to those for a putatively selected class, generating an estimate of the fraction of substitutions that have been fixed by positive selection as opposed to drift. More recent methods for inferring the distribution of fitness effects of new mutations also rely on a neutrally evolving class of site, especially to correct for the effects of past population changes (Keightley and Eyre-Walker 2007; Boyko et al. 2008; Eyre-Walker and Keightley 2009; Schneider et al. 2011; Galtier 2016; Kim et al. 2017; Tataru et al. 2017).

In a number of species of *Drosophila*, there is evidence for the functional significance of a large fraction of the genome, including the action of both purifying and positive selection on intronic and intergenic sites (Andolfatto 2005; Haddrill et al. 2005; Halligan and Keightley 2006; Haddrill et al. 2008; de Procé et al. 2012). Synonymous changes may also be subject to weak selection for preferred codons, which can affect allele frequencies in species with sufficiently large population sizes for such selection to be effective, including *Drosophila melanogaster* and *D. simulans* (Begun 2001; Vicario et al. 2007; Jackson et al. 2017; Machado et al. 2020).

A candidate for a class of neutral nucleotide site in *Drosophila* is the 8-30 basepair region from the 5’ end of introns shorter than 66bp, but ≥ 23bp after removing splice sites (referred to here as SI sites), which have been shown to have the highest between-species divergence and within-population diversity compared to other regions of the genome (Halligan and Keightley 2006; Parsch et al. 2010). These patterns are suggestive of a low level of purifying selection, and consequently a lack of functional importance. This makes them a good candidate for a neutral comparator of the type required by the methods mentioned above. For example, short introns have been used to infer strong purifying selection at 4-fold degenerate sites in *D. melanogaster* (Lawrie et al. 2013; Machado et al. 2020), to fit demographic models to North American *D. melanogaster* in order to determine appropriate parameters for inferring selection from haplotype statistics (Garud et al. 2015), and to quantify population structure in European *D. melanogaster* (Kapun et al. 2020). Sites outside the 8-30bp region but within short introns are probably more constrained because they are functionally important for mRNA splicing (Green 1986; Mount et al. 1992; Kennedy and Berget 1997; Halligan and Keightley 2006).

If SIs do indeed evolve in the absence of selective constraint, they provide an opportunity to investigate processes that affect the composition of genomes other than natural selection. Subsequent studies of evolution at SI sites in *Drosophila* have provided evidence for context-dependent mutational patterns (Clemente and Vogl 2012) and the possible action of GC-biased gene conversion (gBGC) (Jackson et al. 2017). Evidence for gBGC in *Drosophila* genomes is equivocal, with some suggestion that it operates on the X chromosome in *D. simulans* (Haddrill and Charlesworth 2008) and *D. americana* (de Procé et al. 2012), and on both autosomes and the X chromosomes in *D. simulans* and *D. melanogaster* (Jackson et al. 2017), while other studies have found little or no evidence for it (Clemente and Vogl 2012; Comeron et al. 2012; Campos et al. 2013; Robinson et al. 2014).

Direct observations of spontaneous mutations in *D. melanogaster*, as well as analyses of rare segregating polymorphisms, show that mutation is biased towards GC to AT basepair substitutions (Assaf et al. 2017), and population genetic studies have suggested that the extent of this bias has increased at some point in the evolutionary past (Kern and Begun 2005; Zeng and Charlesworth 2010; Clemente and Vogl 2012). Laboratory studies of mutation are limited in power because mutations are rare – with a mutation rate of 5 x 10^-9^ per basepair (Assaf et al. 2017) we expect 0.7 mutations per haploid genome per generation in a genome containing 140 million basepairs. But an examination of the population genetics of natural populations provides the opportunity to integrate evidence from over long evolutionary timescales. In combination with information from sections of the genome that are free from directional evolutionary processes, this procedure should provide further insights into the nature of spontaneous mutations in *Drosophila* and the forces affecting mutations with little or no effects on fitness.

To investigate possible non-selective directional evolutionary processes in *Drosophila*, we have investigated evolution at short intron (SI) sites, using polymorphism data from populations from the putative ancestral ranges of *D. melanogaster* and *D. simulans*, as well as data on between-species divergence. Our study refines the analyses of Jackson *et al.* (2017), which primarily focussed on four-fold degenerate sites, since their analyses of SI sites were hampered by insufficient amounts of data and a poorer quality annotation of the *D. simulans* genome than the one used here. We show that a subset of SI sites are probably subject to directional evolutionary pressures, with GC alleles being favoured over AT alleles at SI sites with the highest GC contents, suggesting the action of GC-biased gene conversion in both species. This has implications for the use of short introns as a neutrally-evolving reference in population genetics, and also sheds light on the dynamics of genome evolution in *Drosophila*. The study also provides further evidence for the existence of a strong GC to AT mutational bias in Drosophila. Its magnitude appears to be independent of the GC content of short introns, and has apparently increased along the *D. melanogaster* lineage following its divergence from the common ancestor of *D. melanogaster and D. simulans.*

## Materials and Methods

### Sequence data from *D. simulans* and *D. melanogaster*

We have analysed a previously published population sample of 21 lines of *Drosophila simulans*, derived from the putatively ancestral Madagascan population (the MD lines of Jackson et al. 2017). The sampling, maintenance, sequencing and variant-calling procedures for these lines were fully described in Rogers *et al.* (2014), Jackson *et al.* (2017) and Becher *et al.* (2020). Briefly, publicly available raw read data in FASTQ format for these 21 lines were downloaded from the European Nucleotide Archive and mapped to version 2.02 of the *D. simulans* reference genome (FlyBase release 2017_04) using BWA MEM (Li and Durbin 2009). We sorted, merged and marked duplicates on the resulting BAM files using Picard Tools version 2.8.3 (https://broadinstitute.github.io/picard/). Variants were called for each line individually using the HaplotypeCaller tool from GATK version 3.7 (McKenna et al. 2010) with the options –emitRefConfidence BP_RESOLUTION and –max-alternate-alleles 2. VCF files containing all 21 lines were generated from the output of HaplotypeCaller using the GATK tool GenotypeGVCFs. We treated sites that remained heterozygous within samples after inbreeding as follows: at each heterozygous site within a sample, one allele was chosen as the haploid genotype call at that site with a probability proportional to its coverage in the sample. The alternative allele was discarded (Jackson et al. 2017).

We also downloaded publicly available sequence data for 197 lines of *D. melanogaster* sampled from Zambia (ZI lines) from the Drosophila Genome Nexus (DGN) (https://www.johnpool.net/genomes.html), and converted these data to FASTA format using a custom shell script. Using the information reported in the supplement to Lack et al. (2015) we retained 69 ZI lines that showed no evidence of admixture with European populations. The sampling, sequencing and variant-calling procedures, and the procedure for defining admixture tracts for these ZI lines were described fully in Pool et al. (2012) and Lack et al. (2015). The ZI sample is maximally diverse and minimally affected by cosmopolitan admixture among the populations in the DGN (Lack et al. 2015), and also provides the largest sample of African *D. melanogaster* genomes.

### Between-species alignments

We used the multispecies alignment between *D. melanogaster, D. simulans* and *D. yakuba* from Zeng *et al.* (2019). Briefly, multispecies alignment was performed between the reference genomes of *D. simulans* (v2.02), *D. melanogaster* (v5.57), and *D. yakuba* (v1.05) using the MULTI-Z pipeline described by Barton and Zeng (2018). Reference genomes were downloaded from FlyBase, and repeat regions were soft-masked using RepeatMasker (http://www.repeatmasker.org/) with the default database for *Drosophila.* Pairwise alignments were generated between *D. melanogaster* and *D. simulans*, and between *D. melanogaster* and *D. yakuba* using LASTZ (Harris 2007), which were chained and netted using axtChain and chainNet (Kent et al. 2003). Single coverage was generated using single_cov2.v11 from the MULTIZ package (Blanchette et al. 2004) and the pairwise alignments were aligned with MULTIZ to create a three-way multiple alignment.

### Defining short intronic sites

To define short intronic (SI) sites, we first carried out the following procedure separately for each of *D. melanogaster* and *D. simulans*. We used the information in the header lines of the FlyBase FASTA file of introns for version 2.02 (5.57) of the *D. simulans* (*D. melanogaster*) reference genome to extract coordinates of the 8–30 bp region of introns that were ≤ 65 bp in length, after checking that this region did not overlap with an exon, an intron of length more than 65 bp, or the non-8–30 bp portion of an intron of length ≤ 65 bp, using information from the gff format annotation of the *D. simulans* (*D. melanogaster*) reference genome version 2.02 (5.57) downloaded from FlyBase.

Using the resulting short intronic positions in each species and the whole genome alignment described above, we defined a set of homologous sites that were annotated as short intronic sites in both *D. simulans* and *D. melanogaster*, using the script non_ref_intersect.py from the WGAbed package (https://henryjuho.github.io/WGAbed/) and the bedtools subroutine intersectBed (Quinlan and Hall 2010), as well as additional custom shell and Python scripts. We generated an alignment for each *D. melanogaster* short intron region that overlapped with a *D. simulans* short intron region, yielding polymorphism data for the ZI and MD lines, as well as the corresponding sequences from each of the *D. melanogaster* v5.57, *D. simulans* v2.02 and *D. yakuba* v1.3 reference sequences.

At this stage, we retained SI sites only if the following additional conditions were met: they were located on an autosome in both *D. melanogaster* and *D. simulans*; there were no missing alleles in any of the three reference sequences; they were not soft-masked as repetitive in any of the three reference sequences; they did not overlap with an indel in the *D. simulans* variant callset; QUAL >= 30 in the *D. simulans* variant callset; they did not lie in a non-crossover region in either the *D. melanogaster* genome (as defined in Campos et al. 2012) or in the *D. simulans* genome (as defined in Becher et al. 2020). In total, we retained 167,147 autosomal SI sites for divergence-based analyses. For polymorphism-based analyses, such as using derived allele frequencies or site frequency spectra (see below), we further excluded sites with any missing polymorphism data in the population under consideration. We retained 163,998 and 145,747 sites for polymorphism-based analyses of the MD lines and the 69 ZI lines, respectively.

For the purposes of comparing regions with different GC contents, we assigned SI sites to bins of equal numbers of introns, according to their GC content. We calculated the GC contents of SI sites from the full 8-30bp region of the short intron under consideration in the reference sequences for *D. melanogaster* and *D. simulans.* Then we assigned SI sites to bins in three different ways. Firstly, we took the mean of the GC content values for each homologous pair of introns and applied this single value when grouping both species’ SIs. This results in the same set of homologous sites in each bin for analyses of divergence and polymorphism in both lineages. Secondly, we grouped introns into species-specific bins, for *D. melanogaster* by using the GC content calculated from the *D. melanogaster* reference sequence, and for *D. simulans* by using the GC content calculated from the *D. simulans* reference sequence. This means that homologous sites may be assigned to different bins in analyses of the *D. simulans* lineage from those in analyses of the *D. melanogaster* lineage, but ensures a perfect relationship between within-species GC content and bin. Below, we refer to these two binning strategies as ‘mean’ and ‘species’, respectively. The two GC contents are closely correlated, as is expected given the slow evolution of GC content (Supplementary Figure S1), but their grouping of SI sites sometimes resulted in contrasting conclusions about patterns of evolution, which we discuss below. Thirdly, we grouped SI sites into bins by the difference in GC content between *D. melanogaster* and *D. simulans* per short intron. We argue below that binning by the mean criterion for GC content is appropriate for analyses of substitution patterns, whereas binning by the species criterion is more appropriate for polymorphism analyses. Binning by the between-species difference in GC content sheds light on processes in regions of the genome that have diverged in GC content between the two species.

To obtain confidence intervals (CIs) around point estimates of statistics, we bootstrapped by sampling introns with replacement 1000 times until the bootstrapped sample was the same size as the observed sample. For each bootstrap sample, we recalculated the statistic of interest. We used the 2.5% and 97.5% quantiles of the resulting distribution as the upper and lower bounds of the 95% CI (Efron 1979).

### Analyses of between-species divergence

We used the GTR-NHb model of base substitution modified to generate sub-optimal ancestral sequences, as implemented in the baseml program of PAML version 4.8 (Yang 2007), in order to reconstruct the base content of the *melanogaster*-*simulans* (*ms*) ancestor, and counted substitutions along lineages according to the Expected Markov Counting method of Matsumoto *et al.* (2015). This method should be more accurate than maximum parsimony or use of a single best reconstruction under complex patterns of base substitution, which are likely apply to *Drosophila* (Matsumoto et al. 2015). We checked that the GTR-NHb fitted the data better than the stationary GTR model, also implemented in PAML, using likelihood ratio tests – this was true in all cases. For each bin, we ran baseml ten times and manually checked for convergence by examining the likelihood output of the model. In the results presented below, we refer to G and C alleles as strong (*S*) and to A and T alleles as weak (*W*). We categorised the number of substitutions from the *ms* ancestor to the extant *simulans* sequence or to the extant *melanogaster* sequence into the following classes: the total number of substitutions from strong to weak alleles, *N_S>W_*; the total number of substitutions from weak to strong alleles, *N_W_>_S_*; and the total number of substitutions from strong to strong alleles or from weak to weak alleles, *N_neu_*. We denote the number of GC sites in the ancestral sequence by *L_GC_* and the number of AT sites in the ancestral sequence by *L_AT_* We define the substitution rate from strong to weak alleles as *r_S>W_* = *N_S>W_*/ *L_GC_*, and the substitution rate from weak to strong alleles as *r_W>S_* = *N_W>S_*/ *L_AT_* We obtained the expected numbers of substitutions and the predicted ancestral base content by parsing the output of PAML, using custom scripts in R (R Core Team 2018).

### Analyses of polymorphism data

We divided SI sites into the same sets of five bins as used for the divergence-based analyses. For each population, we excluded sites with missing data in the polymorphism sample as well as segregating sites of more than two alleles, and then used est-sfs v2.03 (Keightley and Jackson 2018) to calculate the probability of the major allele being ancestral for each segregating site. We used the Kimura 2-parameter model of base substitution and two outgroups (*D. yakuba* and one of either *D. melanogaster* or *D. simulans*, depending on the species to which the polymorphism data belonged) to run est-sfs, and carried out 10 maximum likelihood searches for each bin to check for convergence. Using the results, we constructed separate unfolded site-frequency spectra (SFSs) for segregating *S* >*W*, *W>S* and neutral (*W* >*W* or *S* > *S*) mutations. We used these SFSs to calculate the mean derived allele frequency (DAF) for each class of change, and as an input for the method of Glémin *et al.* (2015) for estimating the mutation and selection parameters.

This method uses the three unfolded SFSs for segregating sites described above to estimate *γ* and *κ*, where *γ* = 4*N_e_S* is the scaled strength of selection for GC (*S*) alleles, and *κ* is the mutational bias parameter *u/v*. Here, *S* is the selection coefficient against heterozygotes for W and S alleles (semi-dominance is assumed), *u* is the mutation rate from *S* to *W*, and *v* is the mutation rate from *W* to *S*. This method uses the three unfolded SFSs described above to disentangle *γ* and *κ*. The method is capable of taking into account polarization errors, which can lead to upwardly biased estimates of *γ* (Hernandez et al. 2007), by incorporating them into the model and estimating them jointly with the parameters of interest. It is also capable of correcting for demographic effects, by introducing nuisance parameters to correct for distortions in the SFS due to demography (following Eyre-Walker et al. 2006). We estimated *γ* and *κ* using the R code provided in the supplement of Glémin et al. (2015). We refer to the models using this method with the same notation as in Glémin et al. (2015). These are model M0, with *γ* = 0, and no correction for polarization errors; M1, with *γ* ≠ 0 and no correction for polarization errors; and M0* and M1*, which are the equivalent models after correcting for polarization errors. We compared the different models using likelihood ratio tests.

### Computational methods

This work made use of GNU parallel (Tange 2011).

### Data Availability

All the code required to replicate the analyses presented here is available on Github (https://github.com/benjamincjackson/dros_gBGC), including the final dataset of homologous short intronic sites in plain text format.

## Results

### Summary of polymorphism and divergence results

In agreement with previous analyses (Jackson et al. 2017), the population of *D. simulans* from its putatively ancestral range in Madagascar is more diverse than either population of *D. melanogaster* – nucleotide diversity (*π*) at SI sites is 0.037 for the MD sample and 0.016 for the ZI sample (Table 1). The site frequency spectrum is more skewed towards rare variants in *D. simulans* than in *D. melanogaster*, as shown by the larger absolute values of Tajima’s *D* and the proportion of singletons in the MD sample, compared to ZI (Table 1). The ratios of the proportions of singletons to their expected values under neutrality are 1.83 and 1.48 for MD and ZI, respectively. Parsch et al. (2010) reported the divergence between *D. melanogaster* and *D. simulans* at the 8-30bp region of introns < 66bp long to be 0.123, with the divergence along the *D. melanogaster* lineage equal to 0.064. These values are nearly identical to those for our dataset with all sites concatenated: 0.124 (0.121-0.126) for the *D. melanogaster* and *D. simulans* net divergence, 0.0655 (0.0637-0.0674) for the *D. melanogaster* branch, and 0.0582 (0.0567, 0.0598) for the *D. simulans* branch (the brackets indicate the confidence intervals for the means).

**Table 1.** Polymorphism statistics for SI sites (means with their 95% CIs)

| Population | Nucleotide diversity ($\pi$) | Watterson's $\theta$ | Tajima's $D$ | Proportion of<br>singletons |
| --- | --- | --- | --- | --- |
| MD21 | 0.0371 (0.0365, 0.0377) | 0.0513 (0.0507, 0.0521) | -1.16 (-1.133, -1.183) | 0.513 (0.506, 0.519) |
| ZI69 | 0.0164 (0.0159, 0.0168) | 0.0195 (0.0191, 0.0199) | -0.566 (-0.517, -0.616) | 0.308 (0.301, 0.317) |

### Testing for fixation bias

If base composition is at statistical equilibrium under mutation, drift and selection over the period of time covered by an evolutionary lineage, there should be equal numbers of substitutions from *S* (G or C) to *W* (A or T) alleles and from *W* to *S* alleles (Bulmer 1991; Akashi 1995; Charlesworth and Charlesworth 2010 p.272). Using the same methodology as here, we previously reported a slight overall AT-bias in substitutions in *D. simulans* autosomal short introns (χ^2^ = 5.55, df = 1, *P* = 0.019) (see Table 2 in Jackson et al. 2017). The results obtained from our new dataset suggests an equilibrium base composition in *D. simulans* autosomal SIs when we concatenate all SI sites. *N_W>S_* for all SI sites combined in *D. simulans* was 3500, and *N_S>W_* was 3584, which do not differ significantly from a ratio of 1:1 (χ^2^ = 0.99, df = 1, *P =* 0.32).

After binning SIs by their mean GC content across species, there was no obvious relationship between GC content and the ratio *N_W>S_*/*N_S>W_*. The 95% confidence intervals obtained by bootstrapping overlap unity for all bins (Figure 1A). This suggests that aggregating sites with different GC contents does not mask any substitution patterns that are specific to sequence context. In agreement with previous results (Akashi et al. 2006; Jackson et al. 2017), *D. melanogaster* shows an overall bias towards AT-fixation for all sites combined (*N_W>S_* = 3478, *N_S>W_* = 4760, χ^2^ = 199, df = 1, *P* < 0.001). When *D. melanogaster* SI sites are binned by mean GC content, all of the bins exhibit an AT fixation-bias, which increases with increasing GC content (Figure 1B). This implies that *D. melanogaster* has experienced a change in the forces acting on GC content, such that its GC content is currently not at equilibrium; this could either be a change in mutational bias towards GC>AT mutations, or a reduction in the scaled intensity of a selective force or biased gene conversion favouring GC, possibly reflecting a reduction in effective population size. In contrast, *D. simulans* appears to be approximately in equilibrium.

**Figure 1.**
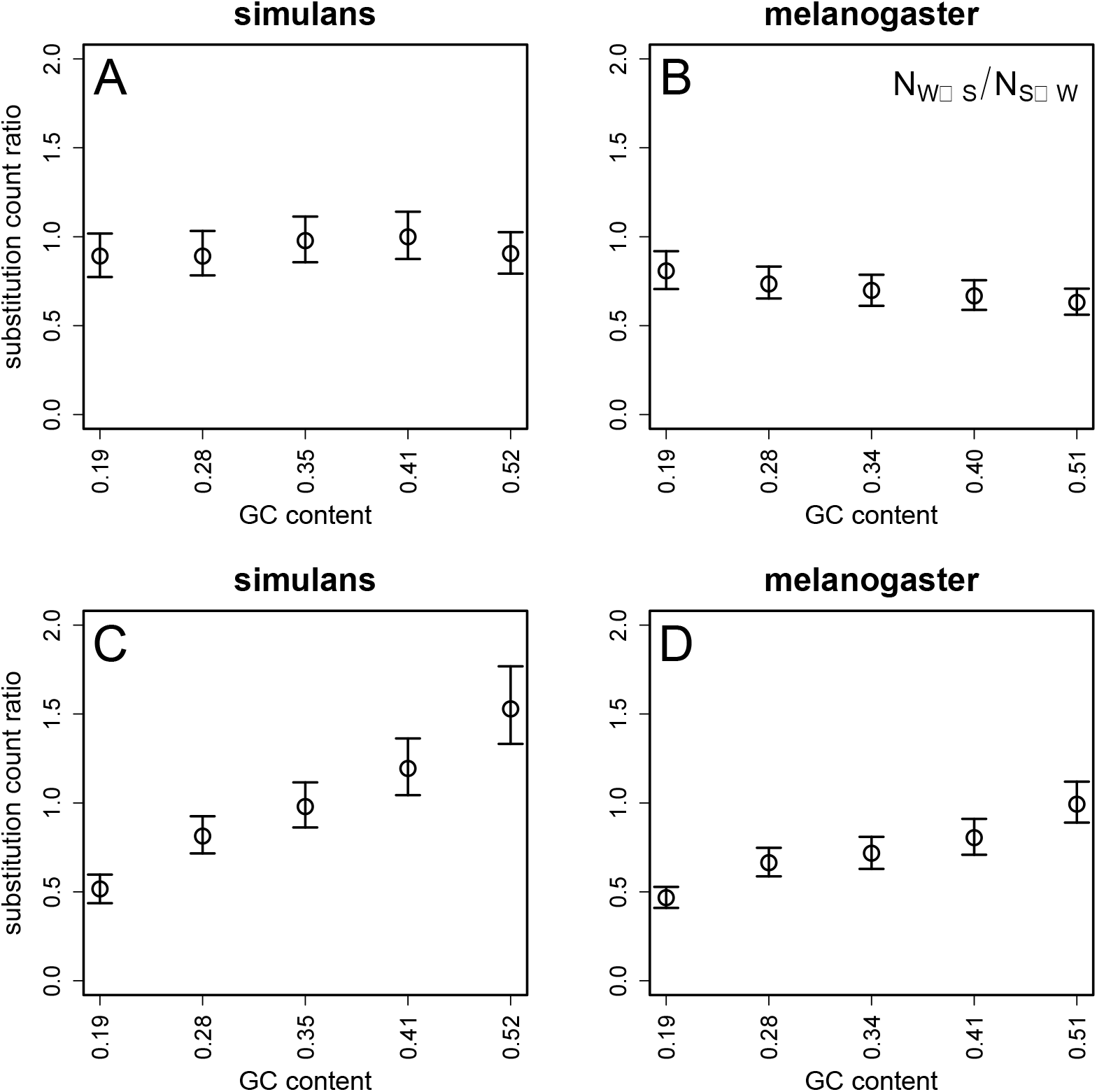
The ratios of substitution counts, plotted against the GC content of SI bins. A substitution count ratio of *N_W>S_*/*N_S>W_* = 1 implies equilibrium base composition. Ratios were calculated for the *D. simulans* lineage (panels A; C) and the *D. melanogaster* lineage (B; D). Sites were binned by the mean GC content of homologous introns in *D. melanogaster* and *D. simulans* (A; B) or by the species-specific GC content (C; D). Error bars represent 95% CIs from 1000 bootstraps of the data in each bin.

On the other hand, when we used the species-specific GC content for binning SIs, in both *D. simulans* and *D. melanogaster* the ratio *N_W>S_*/*N_S>W_* increases with increasing GC content (Figure 1C, D) – i.e. there have been proportionally more changes from AT to GC than from GC to AT in more GC-rich regions. In *D. simulans*, only the middle bin appears to be at equilibrium according to this analysis. The lowest two GC content bins exhibit an excess of *S>W* substitutions, and the highest two GC content bins exhibit an excess of *W>S* substitutions (Figure 1C). In *D. melanogaster*, the bottom four GC bins exhibit an excess of *S>W* substitutions, and the highest GC content bin might be at equilibrium (Figure 1D). There are, however, strong reasons for thinking that this method of binning produces biases in inferences concerning the relationship between substitution patterns and GC content. Consider, for example, the bin with the lowest GC content in a given species. With a substitution rate of around 0.06 along its lineage back to its common ancestor with the other species, the expected number of changes within an SI along the lineage is of the order of 0.06 x 23 = 1.4, or less. The chance that both of these substitutions are both *W>S* is thus very high. If SIs that have the lowest GC content are chosen according to the species-specific GC content, these are automatically enriched for excess of *W>S* changes as opposed to *S>W* changes. The converse applies to SIs chosen for a low GC content. No such selection bias is expected if SIs are chosen on the basis of the mean of their GC contents for the two species. For this reason, the estimates based on the mean bins are more likely to represent the underlying evolutionary process.

### Analyses of polymorphism data

The polymorphism data were used to investigate the parameters of mutation and selection acting on GC versus AT variants. In this case, binning SIs by the species-specific GC content is probably more appropriate than using the mean values across species, since it provides a better reflection of the sequence composition over the comparatively short time-scale experienced by currently segregating variants.

We examined the nature of the forces acting on polymorphic variants in two ways. First, for each bin, we calculated the mean derived variant frequency for different classes of mutations at segregating sites. These classes involved either GC to AT variants (*DAF_S>W_*), AT to GC (*DAF_W>S_*), or GC to CG or AT to TA (*DAF_neu_*). These statistics should shed light on the processes of interest in *Drosophila* genome evolution as follows. If the mutational process has shifted towards a greater GG>AT bias within the last 4*N_e_* generations (the time frame relevant to polymorphism data), as has previously been suggested for *D. melanogaster* (Akashi et al. 2006; Jackson et al. 2017), we expect *DAF_S>W_* < *DAF_W>S,_* because mutations from GC to AT should be younger on average, even under neutrality. Furthermore, if such a change in mutational bias were genome wide, we do not expect a relationship between DAF and GC content. In contrast, if gBGC or a selective force favouring GC is in operation, we expect to have *DAF_W>S_ > *DAF_neu_* > DAF_S>W_* (Jackson et al. 2017). In addition, if the strength of such a force varies across the genome and has influenced its GC content, we expect this relationship be stronger in regions of higher GC content – that is, *DAF_S>W_* should be negatively related to GC content and *DAF_W>S_* should be positively related to GC content. This second pattern, suggestive of a force favouring GC, is indeed what we observe in *D. simulans*, for both the mean and species-specific method of binning SIs by GC content, with a clearer pattern for the species-specific bins than the mean bins (Figure 2A and 2C).

**Figure 2.**
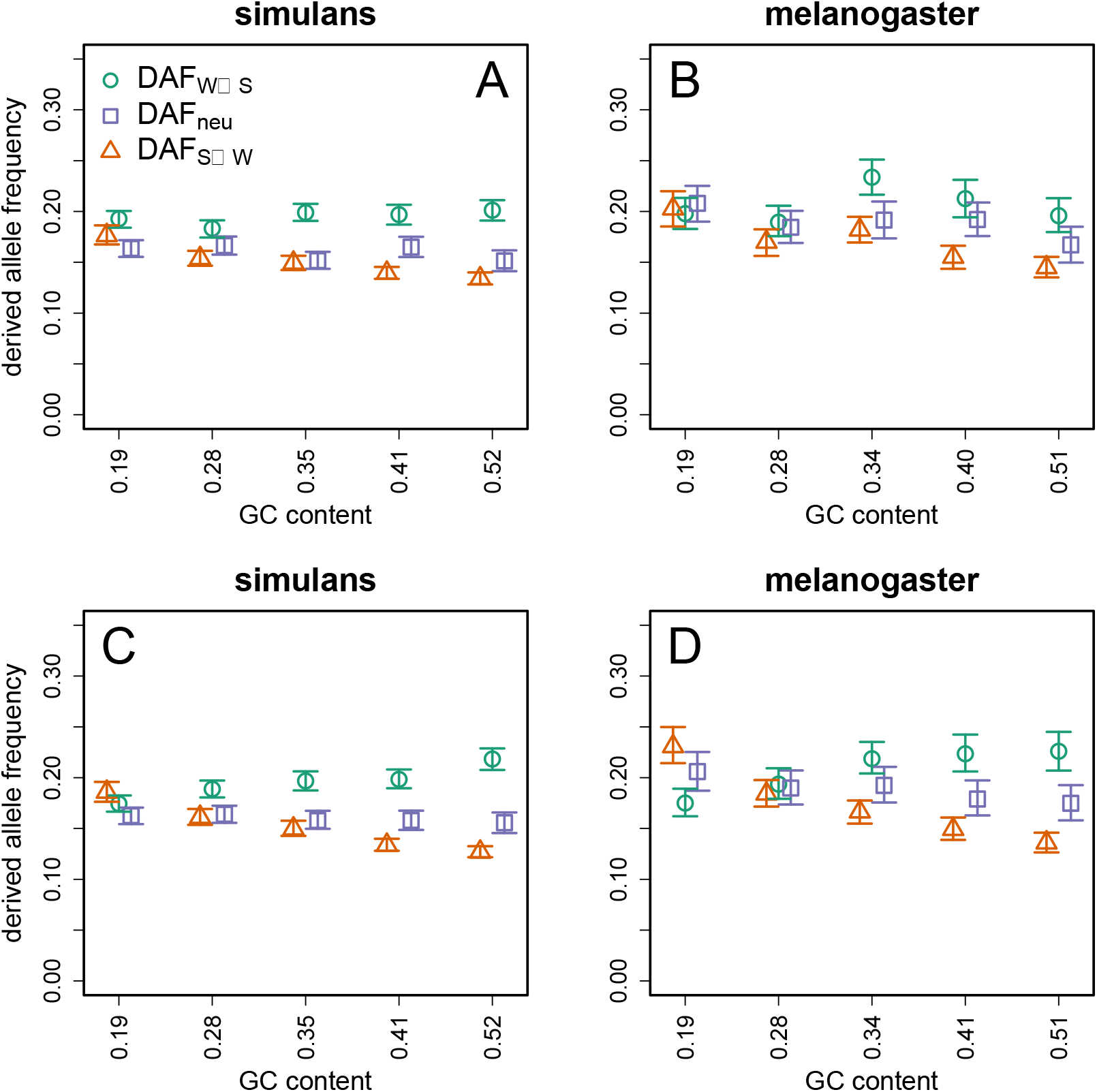
Derived allele frequencies (DAF) for different types of mutations, plotted against the GC content of SI bins. DAF was calculated for the *D. simulans* MD population (panels A; C) and the *D. melanogaster* ZI population (B; D). Sites were binned by the average GC content of homologous introns in *D. melanogaster* and *D. simulans* (A; B) or by species-specific GC content (C; D). Error bars represent 95% CIs from 1000 bootstraps of the data in each bin.

For the Zambian sample of *D. melanogaster*, we observe the pattern supporting a GC favouring force in the top three highest GC content bins when we bin introns using the species-specific method (Figure 2D), but the relationships are less clear when the mean bins are used (Figure 2B). In the case of the mean bins, the three highest GC content bins exhibit *DAF_W>S_* > *DAF_S>W_*; although *DAF_S>W_* seems to decrease with decreasing GC content across all five bins, there is no obvious relationship between *DAF_W>S_* and GC content (Figure 2B).

One possible explanation is that these differences between the mean and species bins reflect the fact that the GC content of the mean bins reflects the ancient GC contents of the SIs involved. If, for example, there has been a shift towards a weaker force favouring GC in both lineages, but whose strength is nevertheless still correlated with GC content (as indicated by the analysis of *γ* below), the mean bins will on average be associated with the past GC content of the bins, so that there will be a less clear relationship of their derived variant frequencies to GC content than for the species bins. The polymorphism results for the species bins would then provide a more reliable picture than those from the mean bins.

Overall, these results suggest the presence of a GC-favouring force in *D. simulans* and, probably to a lesser extent, in *D. melanogaster*. In order to quantify this force, we used the method of Glémin *et al.* (2015) to calculate *γ* = *4N_e_s*, the scaled strength of selection or biased gene conversion favouring GC alleles. In no cases did models correcting for polarisation errors fit the data better than the equivalent model without corrections. This may be because the method we used to polarise segregating sites is less prone to mis-inference than methods such as maximum parsimony (Keightley and Jackson 2018). Both sets of models returned very similar values of *γ* within each species (Figure 3). Below, we report the results from models M0 and M1, which do not correct for polarization error (see Methods).

**Figure 3.**
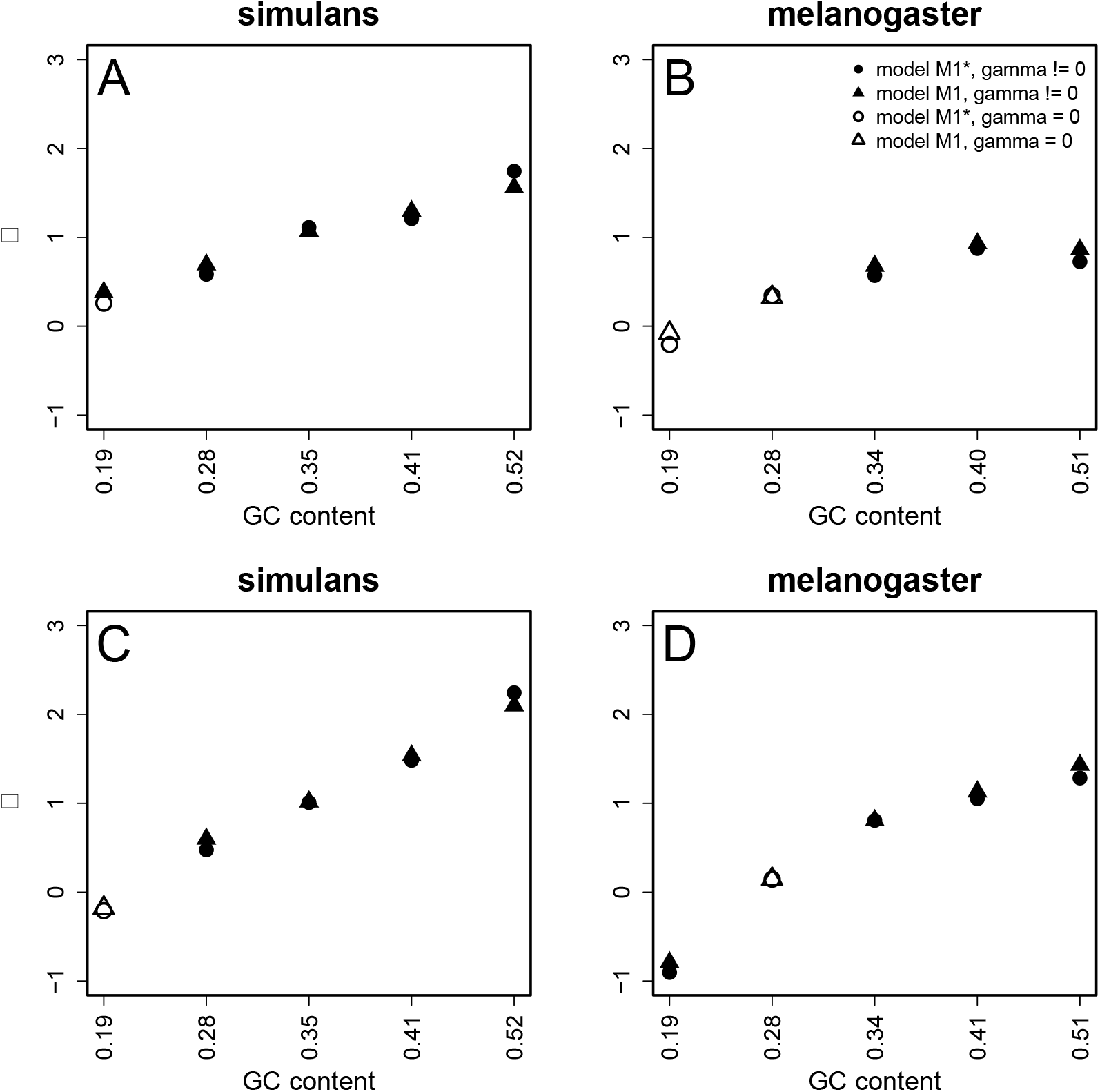
Estimates of the strength of selection in favour of GC alleles (y = 4*N_e_s*), plotted against the GC content of SI bins. Open symbols indicate values which are not significantly different from zero. Circular symbols denote models incorporating parameters that correct for polarisation error. Triangular symbols denote models not incorporating parameters that correct for polarisation error. *γ* was calculated for *D. simulans* (panels A; C) and *D. melanogaster* (panels B; D). Sites were binned by the mean GC content of homologous introns in *D. melanogaster* and *D. simulans* (panels A; B) or by the species-specific GC content (panels C; D).

In *D. simulans*, there is little evidence for a force favouring GC alleles in the lowest GC content bin, with the exception of model M1 in the lowest mean bin, which yields an estimate of *γ* = 0.38 (Figure 3A). In the remaining four bins, *γ* is significantly greater than zero, and increases monotonically from 0.69 in the second lowest GC bin to 1.56 in the highest GC bin for the mean bins (Figure 3A). For the species bins, the relationship between GC content and gamma is somewhat more pronounced, with *γ* = 0.60 in the second lowest GC bin, rising to *γ* = 2.10 in the highest GC content bin (Figure 3C).

For the mean bins in *D. melanogaster*, the pattern is similar, but the force favouring GC is weaker. Neither of the lowest two GC bins show evidence that GC is favoured (Figure 3B). The top three GC bins show evidence for a force favouring GC of a similar magnitude to each other (0.68, 0.93 and 0.86 for bins 3, 4 and 5, respectively). For the species bins, the lowest bin shows evidence for a force favouring AT, with *γ* = − 0.79 (Figure 3D). The second lowest GC bin shows no evidence that either strong or weak alleles are preferred, and the top three bins show evidence for a force favouring GC. As was found for *D. simulans*, the relationship between GC content and *γ* is more pronounced for the species bins than the mean bins. Similar to the interpretations discussed for the DAF patterns, it seems plausible that the action of gBGC or selection has diverged somewhat between the two lineages, and that grouping introns by their mean GC content across the lineage dampens the signal of this effect.

This method also allows the estimation of *κ*, the mutational bias parameter (Figure 4), which is estimated jointly with *γ* (Glémin et al. 2015). We report values of *κ* from model M1. In *D. melanogaster, κ* seems fairly insensitive to GC content. From the mean bins, values of *κ* for bins 1 to 5 are 3.22, 3.15, 3.14, 3.13 and 2.67, respectively (Figure 4B). From the species bins, the values of *κ* are 3.09, 3.13, 3.30, 3.10 and 2.77 (Figure 4D). These values are close to the estimate derived from a meta-analysis of direct observations of spontaneous mutations in *D. melanogaster* mutation accumulation experiments, which was 3.35 (95% CIs: 3.00-3.76) (Assaf et al. 2017). In *D. simulans, κ* is somewhat lower, and seems to be slightly negatively correlated with GC content. Estimates for the mean bins are 2.81, 2.56, 2.57, 2.48 and 2.38 for bins 1 to 5, respectively (Figure 4), and estimates for the species bins are 2.82, 2.61, 2.50, 2.53 and 2.40 (Figure 4C). This is in agreement with the hypothesis of an increase in the GC to AT mutational bias in the *D. melanogaster* lineage relative to the *D. simulans* lineage, which has been proposed before (Takano-Shimizu 2001; Kern and Begun 2005; Zeng and Charlesworth 2010; Clemente and Vogl 2012).

**Figure 4.**
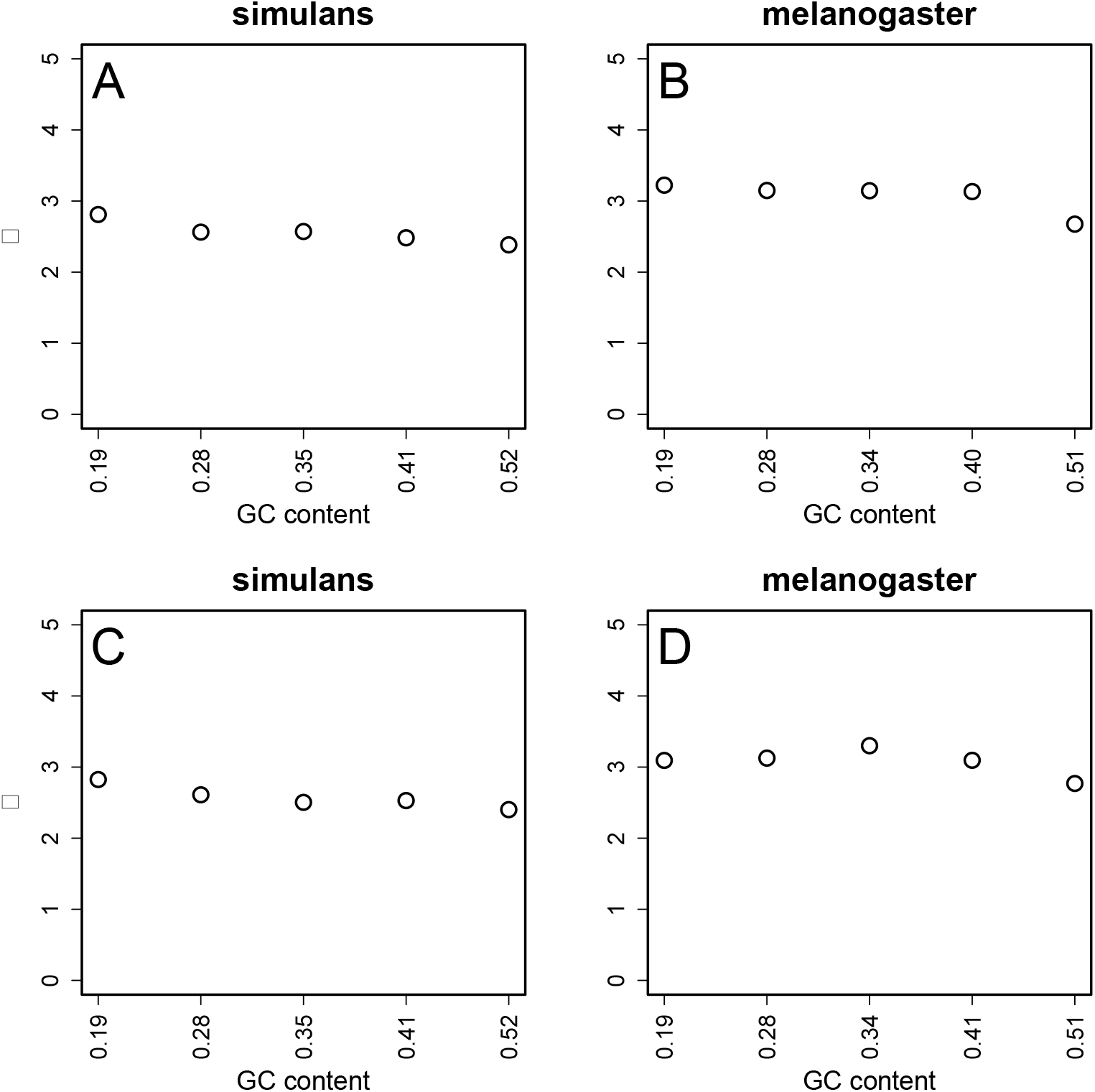
The estimates of the mutational bias parameter, *κ*, calculated using the method of Glémin *et al.* (2015), plotted against the GC content of SI bins. *κ* was estimated from the model M1 (with *γ* != 0 and polarisation errors not corrected for), for both *D. simulans* (panels A; C) and the *D. melanogaster* (B; D). Sites were binned by the mean GC content of homologous introns in *D. melanogaster* and *D. simulans* (A; B) or by the species-specific GC content (C; D).

### Patterns of substitution and their relationship with GC content

If a fraction of SI sites is subject to a weak force favouring GC over AT, this should be reflected in the patterns of substitution rates. As a null hypothesis, we might expect sites in the lowest GC content bins, where there is little evidence for a force favouring GC, to exhibit substitution rates that reflect the mutational bias inferred from polymorphism data and mutation accumulation experiments – under neutrality, substitution rates and mutation rates are equal (Wright 1938; Kimura 1968). For higher GC content bins, where there is evidence for an advantage to GC, we expect a higher rate of substitution of GC alleles relative to the lower GC content bins. To investigate this, we reconstructed ancestral states using PAML (see Materials and Methods for details), and counted the numbers of *S*>*W* and *W*>*S* substitutions along each lineage, in order to estimate the ratio of the two substitution rates, *R* = *r_S>W_*/*r_W>S_*, for both the *D. melanogaster* and *D. simulans* lineages. Note that *R* = *κ* under strict neutrality.

In agreement with this hypothesis, *R* and GC content are negatively correlated in both *D. simulans* and *D. melanogaster* (Figure 5). However, the values of *R* for the lowest mean GC content bins are much higher than the values of *κ* described above. For the lowest mean bins, *R* is equal to 4.56 and 5.03 in *D. simulans* and *D. melanogaster*, respectively (Figure 5A, B). The values of *R* are more extreme for the species than the mean bins (Figure 5C, D) – for the lowest species GC content bins, *R* = 7.51 in *D. simulans* and *R* = 8.19 in *D. melanogaster.* The absolute rates *r_S>W_* and *r_W>S_* are plotted in Supplementary Figure S2. As described above, there are reasons to think that the results for mean bins are more reliable than the species-specific bins, which are likely to be biased towards inferring S>W substitutions. Nevertheless, the fact that *R* >> *κ* for the mean bins with low GC content requires an explanation.

**Figure 5.**
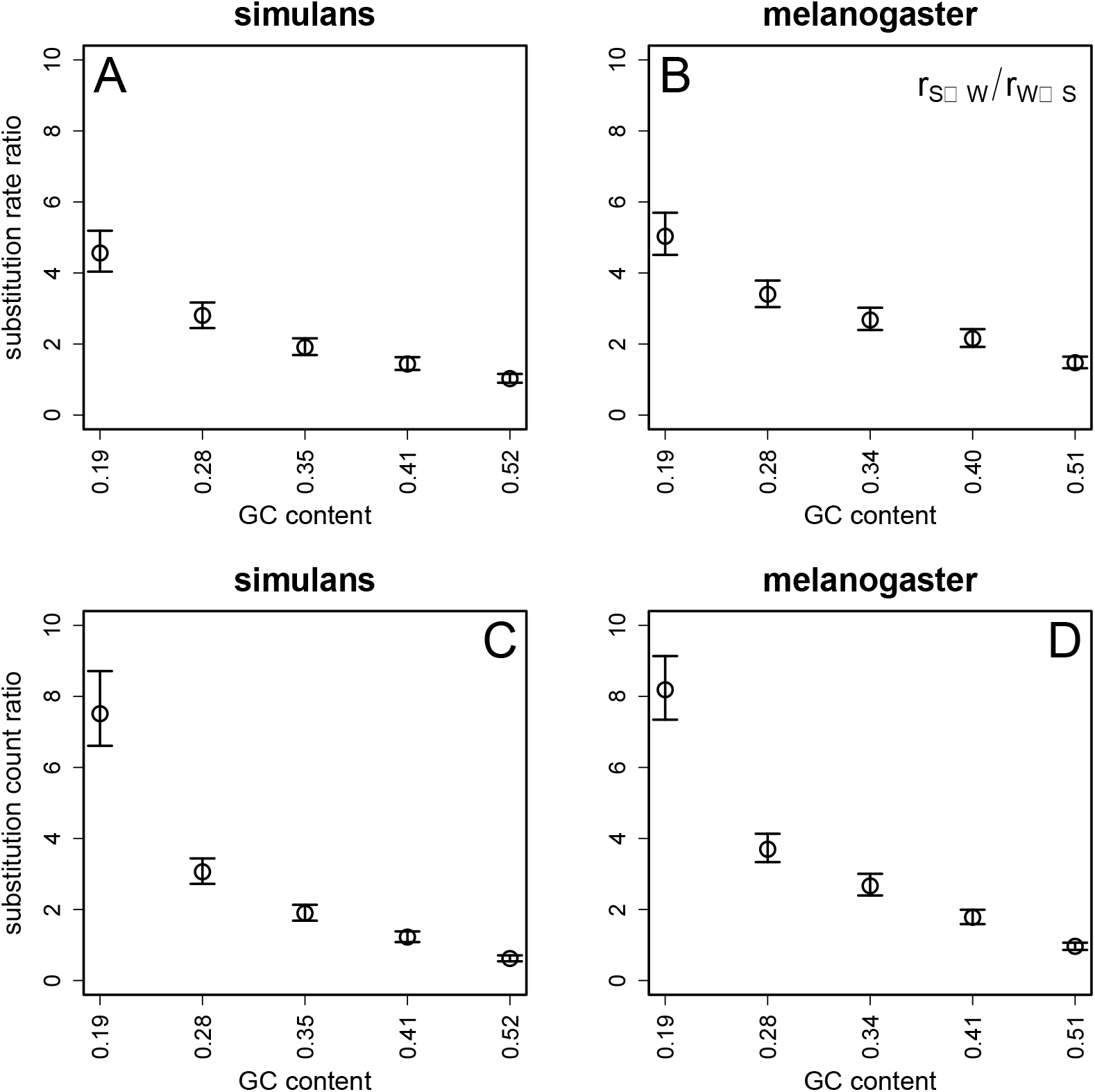
The ratio of substitution rates, *R* = *r_S>W_*/*r_W>S_* plotted against the GC content of SI bins. *R* was calculated for the *D. simulans* lineage (panels A; C) and the *D. melanogaster* lineage (B; D). Sites were binned by the average GC content of homologous introns in *D. melanogaster* and *D. simulans* (A; B) or by species-specific GC content (C; D). Error bars represent 95% CIs from 1000 bootstraps of the data in each bin.

One possibility is that there has been selection in these bins in favour of AT rather than GC along both lineages, consistent with the significantly negative value of *γ* for the lowest GC content species bin in *D. melanogaster* (Figure 3 D). This would also explain why both the present and reconstructed ancestral GC contents of the low GC bins are much lower than the equilibrium GC content predicted on the basis of neutral evolution and the estimated *κ* value of approximately 3 (Figure 4); this is equal to 1/[1 + *κ* exp(-*γ*)], which takes a value of 0.25 with *γ* = 0 (Li 1987; Bulmer 1991; Charlesworth and Charlesworth 2010, p.275). Table S1 shows that, for the lowest mean bin, the GC content for *D. simulans* is 0.195 and the GC content for *D. melanogaster* is 0.192. The GC contents of the ancestral sequence reconstructed by PAML are 0.197 for the lowest *D. simulans* and *D. melanogaster* mean bins. For neutral equilibrium, such a low GC content requires *κ* = 4.13 under neutrality, but a smaller value if there were some selection against GC, e.g. with *κ* = 2.8 (the species value for *D. simulans)*, an equilibrium GC content of 0.20 requires *γ* = – 0.35. From Equation 2 of Jackson et al. (2017), this implies an *R* value of 3.97, which is close to the value estimated for the *D. simulans* lineage. Alternatively, there could have been a shift downwards from an ancestral *κ* value of approximately 4 in both lineages, which would not be detected if it occurred prior to the period of time for which the polymorphism data are relevant (these only provide an estimate of the value of *κ* over the past *4Ne* generations).

Another explanation is that the discrepancy between *R* and *κ* for the low GC bins is an artefact due to alignment errors in AT-rich regions, which might contain more repeat sequences (Bachtrog et al. 1999) and thus be harder to align. The *D. melanogaster* reference sequence is moderately more AT-rich in regions that we soft-masked for repeat content compared to the remainder of the genome (GC content for the masked regions = 39%, GC content for the remainder of the genome = 43%). However, the lowest GC content bins do not exhibit lower alignment scores according to the whole genome alignment MULTI-Z output, which suggests that there are no specific alignment problems for these regions (Supplementary Figure S3). As noted in the methods, we excluded sites masked as repetitive in either species from all of our analyses.

### No effect of polymorphism on patterns of divergence

Because the nucleotide diversity within these species represents an appreciable fraction of divergence between them (Table 1), we repeated our analyses of divergence after removing sites that were polymorphic in either the MD sample or the ZI sample, to approximate a dataset consisting only of fixed differences between *D. simulans* and *D. melanogaster.* Removing polymorphic sites had the effect of considerably reducing the substitution rates for both species, more so for *D. simulans* (compare Supplementary Figures S2 and S5 A, B). It had no effect on the patterns of *R* or the substitution count ratio (Supplementary Figure S5).

## Discussion

Understanding whether genome evolution involves gBGC or a selection pressure acting on the GC content of putatively functionless sequences is important for two reasons. First, it is needed for a complete understanding of the processes affecting the genetic composition of natural populations. Second, we expect it to affect sites that are often used as comparators for detecting other evolutionary processes, such as selection on functionally important sites, changes in population size and mutation. To date, the evidence for a GC-favouring force in *Drosophila* has been ambiguous. It has been claimed to be acting on the X chromosome of *D. simulans* (Haddrill and Charlesworth 2008) and *D. americana* (de Procé et al. 2012), and on both X chromosomes and autosomes in *D. simulans* and *D. melanogaster* (Jackson et al. 2017), while several other studies have failed to find support for it (Clemente and Vogl 2012; Comeron et al. 2012; Campos et al. 2013; Robinson et al. 2014). We have extended our previous work on this topic (Jackson et al. 2017) by focussing exclusively on short intronic sites, and by using a larger polymorphism sample in *D. melanogaster* than before, together with more complete annotation of the *D. simulans* genome. This allowed us to take the intersection of sites annotated as short introns in both species, which in turn allowed a direct comparison of the processes acting at homologous sites.

Overall, the analyses presented above suggest the existence of a GC-favouring force in both *D. simulans* and *D. melanogaster*, whose strength is positively related to the GC content of an intron, and which is on average stronger in *D. simulans*. This makes sense in the context of GC-biased gene conversion (gBGC), which is a recombination-association process whose evolutionarily effective strength is proportional to the product of the rate of change of allele frequency by gene conversion and the effective population size (*N_e_*) (Nagylaki 1983; Charlesworth and Charlesworth 2010, p.529). On the basis of pairwise diversity at SI sites, *N_e_* is substantially higher for the *D. simulans* population compared to the *D. melanogaster* population, assuming that mutation rates are similar for the two species (Table 1 and Figure S6), so that the evolutionarily effective strength of any deterministic force over the recent past should be larger in *D. simulans*, other things being equal. There are only weak relationships between SI site diversity and GC content (Figure S6), with an observed ratio of *π* for the highest versus the lowest species bins of 0.79 for *D. simulans* and 0.84 for *D. melanogaster.* If mutation-selection-drift equilibrium is assumed, and the estimates of *κ* and *γ* for these species bins are inserted into Equation 15 of McVean and Charlesworth (1999), the predicted ratios are 0.93 and 1.47 for *D. simulans* and *D. melanogaster*, respectively. The agreement between the observed and predicted values is reasonably good for *D. simulans*, but there is a large discrepancy for *D. melanogaster*, possibly reflecting a larger departure from base composition equilibrium than in the case of *D. simulans.*

Nevertheless, the presence of a force favouring GC is suggested both by the analyses of polymorphism data, using estimates of the derived allele frequencies of different sorts of mutation (Figure 2) and the site frequency spectrum based estimate of *γ* (Figure 3), as well as by the analyses of substitution rates (Figure 5). There is also evidence for a strong mutational bias in favour of GC>AT mutations, consistent with direct evidence from mutation accumulation data (Assaf et al. 2017). This bias is larger in *D. melanogaster* than *D. simulans*, and the evidence that base composition is close to statistical equilibrium in the latter but not the former suggests that there may have been a shift towards a stronger mutational bias in the *D. melanogaster* lineage, as has previously been suggested on the basis of somewhat weaker evidence (Kern and Begun 2005; Zeng and Charlesworth 2010; Clemente and Vogl 2012; Jackson et al. 2017). The mechanistic basis and evolutionary significance of such a shift are both unclear.

Our conclusions about the existence of a force favouring GC over AT, and its dependence on the GC content of introns appear to be robust to the different ways of binning SI sites. However, the results also suggest that the conclusions about some of the details of the evolution of SI sites in these two species are sensitive to the methods used to aggregate them, which may be subject to various sources of bias. In particular, binning according to the species-specific GC contents of SIs is likely to produce a bias in favour of W>S substitutions, as discussed in the section on testing for fixation bias, so that the results on substitution patterns from bins based on mean GC content are probably more reliable than those from the species bins.

In contrast, there has been some divergence in GC content at a subset of the SI sites that are shared between *D. simulans* and *D. melanogaster*, with *D. simulans* SIs having slightly higher GC contents than the same introns in *D. melanogaster* (see the rows labelled ‘Mean’ in Table S1). Binning short introns by the mean GC content of homologous sites potentially has the effect of masking some of these differences, because it aggregates sites which are subject to different evolutionary pressures. Use of the species-specific binning method is thus likely to provide a more accurate representation of the sequence context than the use of means, which is relevant to estimates of the strength of gBGC or selection from polymorphism data, although these are qualitatively consistent with the patterns from the mean bins.

We explored these differences further by additionally binning sites by the *difference* in GC content between *D. simulans* and *D. melanogaster* (*GC_diff_*), and observed even more extreme patterns than for the species bins (Supplementary Figure S4), with evidence from the polymorphism data for selection against GC in *D. simulans* in bins for which *GC_diff_* is smallest (and negative), and against GC in *D. melanogaster* in bins for which this difference is the largest. Both *R* and *γ*change with increasing *GC_diff_* in opposite directions in the two species (Figure S4). The relationship between *R* and *GCdiff* is probably partly due to the sampling bias described in the section on testing for fixation bias: selecting introns on the basis of *GC_diff_* causes a bias towards an excess of W>S changes as opposed to S>W changes in the *D. simulans* lineage for introns with a low GC content in *D. simulans*, and vice-versa for *D. melanogaster*.

The complementary relation observed for *γ* is more difficult to explain, since the same sets of introns in each bin are being used for the polymorphism analyses in the two species, yet low *GC_diff_* introns have low *γ* values in *D. simulans* and high *γ* values in *D. melanogaster.* This finding implies that shifts in the base composition of a considerable fraction of introns must have occurred along each lineage as a result of changes in *γ* values in opposite directions in the two lineages. This possibly even includes shifts in the direction of the directional force or forces affecting the frequencies of GC versus AT basepairs, given the evidence for negative *γ* in low *GC_diff_* introns in *D. simulans* and in high *GC_diff_* introns in *D. melanogaster* (Figure S4). Such changes would also contribute to the pattern for *R* described above; Equation 2 of Jackson *et al.* (2017) shows that, with *κ* = 3, a *γ* of – 1 results in *R* = 8.89. It is, however, surprising that they have been sufficiently large to produce shifts in GC content of approximately 0.10 over the relatively short evolutionary timescale involved.

The biological mechanisms underlying the GC-favouring force, if it is real, are unclear. Direct experimental evidence for a GC-bias in transmission of alleles to the products of meiosis due to the repair machinery associated with recombination is limited to budding yeast, where the segregation distortion in favour of GC is modest (Mancera et al. 2008; but see Liu et al. 2018) and to mammals, where the distortion is strong at recombination hotspots (Webb et al. 2008; Duret and Galtier 2009; Arbeithuber et al. 2015). Indeed, Liu et al. (2018) failed to detect significantly GC-biased segregation in yeast, *Neurospora*, *Chlamydomonas* and *Arabidopsis*, although population genetics evidence has suggested its existence in yeast (Harrison and Charlesworth 2011) and *Arabidopsis* (Hämälä and Tiffin 2020). To our knowledge, there is no direct experimental evidence for gBGC, or any other form of biased transmission in *Drosophila.* In mammals, it has been hypothesised that the strength of gBGC is an adaptation to counter the high rate of mutation of methylated cytosines (Brown and Jiricny 1987; Duret and Galtier 2009). *Drosophila* has far lower levels of cytosine methylation compared to mammals (Gowher et al. 2000; Capuano et al. 2014), and the mismatch repair machinery of *Drosophila* differs from mammals and other eukaryotes in important ways (Sekelsky 2017). Consequently, it is unclear *a priori* what level of expectation there is for a GC or an AT bias in transmission of alleles in *Drosophila.* Direct observation of the progenitors and products of meiosis in *Drosophila* would be useful for testing the patterns reported here. Given that there is some doubt about the accuracy of ancestral state inference when the standard assumption of neutral evolution is applied, even using the up-to-date methods employed here, we suggest that work extending models of base composition evolution to incorporate weak directional forces (such as gBGC) would also be worthwhile (e.g., Borges et al. 2019).

## Acknowledgements

The authors thank Kai Zeng for helpful comments on the manuscript, and Henry Barton for providing the multiple alignment between *Drosophila* species.

